# A MAP kinase cascade modulates expression of late G1 phase cyclins in budding yeast

**DOI:** 10.64898/2025.12.02.691939

**Authors:** Navid Zebarjadi, Douglas R. Kellogg

## Abstract

Expression of late G1 phase cyclins is the critical molecular event that marks commitment to enter the cell cycle. In budding yeast, late G1 phase cyclins initiate and sustain growth of a new daughter bud, so their expression also marks the start of a new growth phase during the cell cycle. Expression of late G1 phase cyclins is influenced by nutrient availability – cells growing in poor nutrients progress through late G1 phase with lower levels of late G1 phase cyclins. However, little is known about how or why nutrients modulate expression of late G1 phase cyclins. Here, we investigated the signals that control expression of the late G1 phase cyclin Cln2. We discovered that nutrients modulate expression of Cln2 primarily via post-transcriptional mechanisms that influence Cln2 phosphorylation and turnover. Nutrient modulation of Cln2 protein expression requires a TORC2-Pkc1-MAP kinase signaling axis. Furthermore, expression of Cln2 is closely correlated with bud growth and required for bud growth. A model that could explain the data is that nutrients modulate Cln2 expression to ensure that the rate of bud growth is matched to the availability of nutrients that support bud growth.

## Introduction

In all cells, cell growth generates signals that control the cell cycle to ensure that cell division only occurs when an appropriate amount of growth has occurred (Turner et al., 2012; Jorgensen and Tyers, 2004; Kellogg, 2025). The amount of growth required for cell division varies over many orders of magnitude across the tree of life, leading to an extraordinary diversity of cell sizes (Futcher and Kellogg, 2024). In eukaryotic cells, growth occurs throughout the cell cycle and the requirement for growth is imposed at multiple points during the cell cycle (Liu et al., 2024; Cadart et al., 2018; Varsano et al., 2017; Leitao and Kellogg, 2017; Sommer et al., 2021; Anastasia et al., 2012; Jasani et al., 2020).

A particularly important point at which growth controls the cell cycle is at the end of G1 phase, when cells make the decision to enter a new round of cell division. However, the molecular mechanisms that drive cell cycle entry, and how they are controlled by cell growth, are poorly understood in both yeast and mammals (Rubin et al., 2020). The critical molecular event that marks cell cycle entry is expression of late G1 phase cyclin proteins, which bind and activate Cdk1 to initiate cell cycle entry. It has been thought that expression of late G1 phase cyclins is controlled primarily at the transcriptional level (Breeden, 2003). In budding yeast, transcription of late G1 phase cyclins is controlled by the SBF transcription factor, which is bound and inhibited by a transcriptional inhibitor known as Whi5. An early G1 phase cyclin known as Cln3 was thought to activate Cdk1 to phosphorylate and inhibit Whi5, thereby promoting cell cycle entry (Costanzo et al., 2004; de Bruin et al., 2004). A similar set of signaling steps has been thought to drive cell cycle entry in mammalian cells.

More recent findings in both yeast and mammalian cells are inconsistent with the standard transcriptional model for cell cycle entry and suggest that post-transcriptional mechanisms play a major role. For example, it was found that Cln3 is not required for phosphorylation of Whi5 (Bhaduri et al., 2015; Kõivomägi et al., 2021). Rather, Whi5 phosphorylation is likely driven primarily by a positive feedback loop in which the late G1 phase cyclin Cln2 activates Cdk1 to phosphorylate and inactivate Whi5 (Skotheim et al., 2008). In addition, cells that lack both Cln3 and Whi5 show normal cell cycle dependent expression of Cln2, and Cln2 protein expressed from a constitutive heterologous promoter still undergoes cell cycle-dependent changes in protein levels (Brambila et al., 2024). Finally, Cln3 can influence expression of Cln2 in *whi5Δ* cells and in cells that express Cln2 from a heterologous constitutive promoter (Brambila et al., 2024). These observations show that expression of Cln2 is strongly influenced by post-transcriptional mechanisms. Post-transcriptional mechanisms also appear to play an important role in cell cycle entry in mammalian cells (Narasimha et al., 2014). Little is known about these mechanisms.

Another major gap in our understanding of cell cycle entry concerns the mechanisms by which external signals influence cell cycle entry. In both prokaryotes and eukaryotes, external nutrient status strongly influences the amount of growth required for cell cycle progression – cells growing in poor nutrients grow more slowly and require less growth to progress through the cell cycle, which leads to a substantial reduction in cell size (Kellogg and Levin, 2022). In budding yeast, the effects of nutrients on cell cycle progression were first observed at cell cycle entry (Lorincz and Carter, 1979; Hartwell and Unger, 1977; Johnston et al., 1977). These classical studies found that the duration of G1 phase is increased in poor nutrients, while the extent of growth is reduced, leading to cell cycle entry at a reduced size. Analysis of G1 phase cyclin expression in this context led to surprising findings. In both rich and poor nutrients, Cln3 protein levels increase gradually during growth in early G1 phase, and the rise in Cln3 levels appears to be dependent upon growth and correlated with the extent of growth (Sommer et al., 2021). Given that Cln3 is thought to promote cell cycle entry, one might expect to find that expression is increased in poor nutrients to drive earlier cell cycle entry. Paradoxically, however, expression of Cln3 protein is dramatically reduced in poor nutrients (Sommer et al., 2021). Thus, cells in poor nutrients require less Cln3 for cell cycle entry. Previous studies suggest that cells growing in poor nutrients also require lower levels of Cln2 to enter the cell cycle (Schneider et al., 2004). However, a caveat is that these studies analyzed Cln2 protein expression in asynchronously growing cells so it was not possible to discern Cln2 protein levels at cell cycle entry. Expression of Cln2 protein under different nutrient conditions has yet been analyzed in synchronized cells.

How and why is expression of late G1 phase cyclins reduced when cells are growing slowly in poor nutrients? To begin to answer these questions, we analyzed expression of Cln2 protein in cells growing in rich and poor carbon conditions.

## Results

### Expression of Cln2 is correlated with the rate and extent of bud growth

Previous work found that expression of Cln2 protein is reduced in cells growing asynchronously in poor nutrients (Schneider et al., 2004). To further investigate, we carried out new experiments using cells synchronized in early G1 phase to investigate how nutrient availability influences the expression and requirement for late G1 phase cyclins at cell cycle entry. We used centrifugal elutriation to isolate small newborn daughter cells in early G1 phase. Before elutriation, cells were grown in poor carbon medium (YP + 2% glycerol and 2% ethanol) which leads to decreased cell size and an increase in the fraction of cells in G1 phase. Small newborn cells were isolated and released back into poor carbon medium or into rich carbon medium (YP + dextrose). In this context, the cells in rich carbon must undergo a longer interval of growth to meet the increased growth requirement of rich carbon, while cells in poor carbon require less growth and enter the cell cycle sooner. We took samples at 10-minute intervals and assayed cell size and Cln2 protein levels. We also assayed the percentage of cells undergoing bud emergence as a morphological marker of cell cycle entry, and plotted percent bud emergence as a function of cell size to determine cell size at cell cycle entry (**Figures 1A-D**). The cells in poor carbon initiated bud emergence approximately 30 minutes before cells in rich carbon and they grew more slowly and initiated bud emergence at a smaller cell size. In addition, expression of Cln2 was accelerated in poor carbon and peak levels of Cln2 were lower. In both rich and poor carbon, Cln2 was first detected at 5% bud emergence and then increased steadily during bud growth.

**Figure 1.**
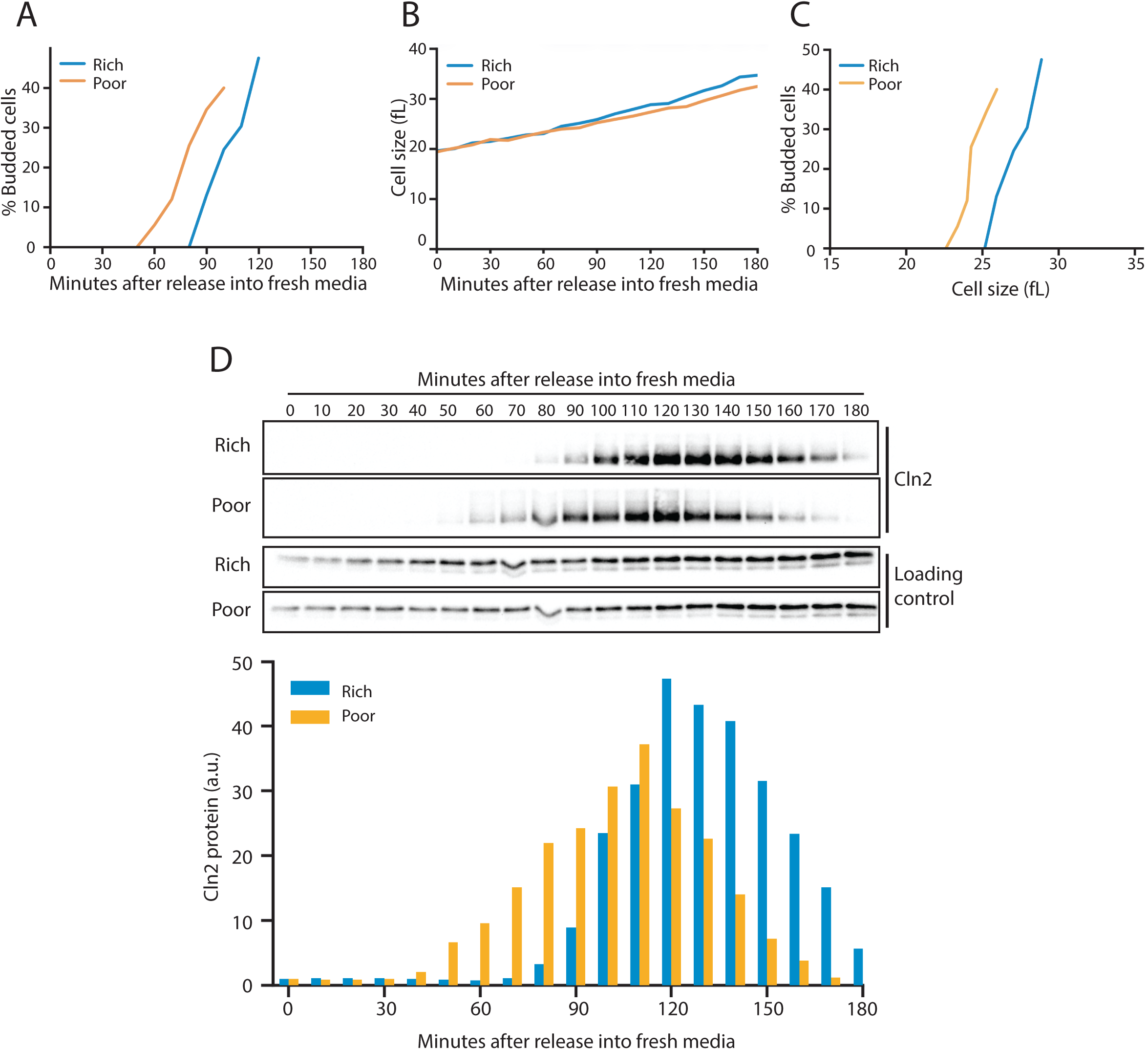
Dynamics of Cln2 protein expression during G1 phase in cells growing in rich or poor carbon. Wildtype cells were grown to midlog phase in poor carbon (YPG/E) and small unbudded cells were isolated by centrifugal elutriation. Cells were released into either rich carbon (YPD) or poor carbon (YPG/E) medium at 25°C and samples were taken at 10 min intervals. All data in the figure are from the same biological replicate. **(A)** The percentage of budded cells plotted as a function of time. **(B)** Median cell size was measured using a Coulter counter and plotted as a function of time. **(C)** The percentage of budded cells as a function of median cell size at each time point. **(D)** The behavior of Cln2-9XMyc was analyzed by western blot. An anti-Nap1 antibody was used as a loading control. The bar graph shows quantification of Cln2-9XMyc protein levels from as a function of time.

These results provide a detailed picture of Cln2 protein expression in G1 phase and provide further confirmation that cells growing slowly in poor carbon require less Cln2 to undergo cell cycle entry. The reduced requirement for Cln2 to pass cell cycle entry in poor nutrients is puzzling. An explanation could come from a previous study that found that Cln2 is not simply required for initiation of cell cycle entry and bud emergence. Rather, Cln2/Cdk1 activity is continuously required for bud growth during the interval of polar bud growth that occurs before mitosis (McCusker et al., 2012, 2007). Thus, rapid inhibition of Cln2/Cdk1 with an analog-sensitive allele of Cdk1 during the polar phase of bud growth leads to an immediate cessation of bud growth. The fact that Cln2/Cdk1 both initiates and drives bud growth suggest a model in which the reduction in Cln2 levels in poor nutrients reflects a need to reduce the rate of bud growth to match the availability of nutrients that are essential for bud growth. A full test of this model is beyond the scope of the present study. Here, we carried out additional experiments to search for the signals the control Cln2 protein levels in response to nutrient availability, with the goal of determining whether regulation of Cln2 is linked to signals that control cell growth.

### Cln2 protein expression responds rapidly to changes in nutrient availability

To further investigate nutrient modulation of Cln2 protein expression, we tested how Cln2 protein responds to an abrupt shift from rich to poor carbon. Cells were grown to log phase in rich carbon and then rapidly shifted to poor carbon and the behavior of Cln2 protein was monitored by western blot. Cln2 underwent immediate hyperphosphorylation within 5 minutes of the shift from rich to poor carbon, which was followed by rapid destruction (**Figure 2A**). We also used northern blotting to monitor Cln2 mRNA levels, which showed that Cln2 mRNA levels decline when cells are shifted to poor carbon (**Figure 2B**). Previous work found that Cln3 shows a similar rapid hyperphosphorylation and destruction in response to a shift from rich to poor carbon (Sommer et al., 2021). The mitotic cyclin Clb2 did not undergo hyperphosphorylation and destruction in response to a shift from rich to poor carbon (**Figure 2C**). These results show that both early and late G1 phase cyclin protein levels are exquisitely sensitive to changes in nutrient availability that strongly influence the rate and extent of growth in G1 phase.

**Figure 2.**
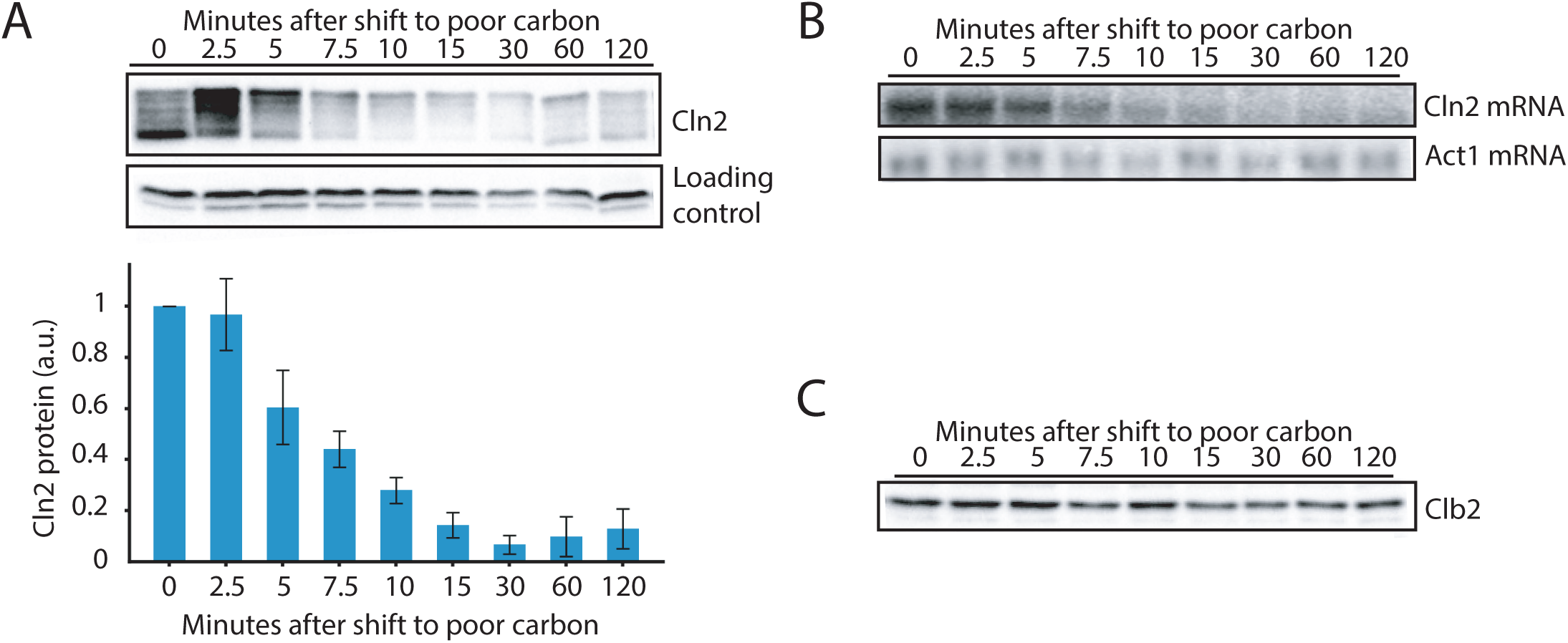
Cln2 protein expression responds rapidly to changes in nutrient availability. **(A)** Rapidly growing cells in rich carbon (YPD) were shifted to poor carbon (YPG/E) at 30°C and the behavior of Cln2-9XMyc was analyzed by western blot. An anti-Nap1 antibody was used as a loading control. **(B)** Cln2 mRNA levels in wild-type cells shifted from rich to poor carbon were analyzed by northern blot. **(C)** Clb2 protein levels in wild type cells after a shift from rich to poor carbon were analyzed by western blot.

### Nutrients modulate expression of late G1 phase cyclin proteins via post-transcriptional mechanisms

It has been thought that expression of late G1 phase cyclins is controlled primarily at the transcriptional level (Breeden, 2003; Costanzo et al., 2004; de Bruin et al., 2004; Kõivomägi et al., 2021). However, recent studies in both yeast and mammals suggest that cell cycle-dependent expression of late G1 phase cyclins is controlled via post-transcriptional mechanisms (Brambila et al., 2024; Narasimha et al., 2014). We therefore tested whether post-transcriptional mechanisms play a role in nutrient-modulation of Cln2 expression. We first tested whether nutrient modulation of Cln2 protein levels requires Cln3 and Whi5, which are thought to be key regulators of *CLN2* transcription. Wild type and *cln3Δ whi5Δ* cells were shifted from rich to poor carbon and the behavior of Cln2 protein was assayed by western blot (**Figure 3A**). Cln3 and Whi5 were not required for nutrient modulation of Cln2 protein levels.

**Figure 3.**
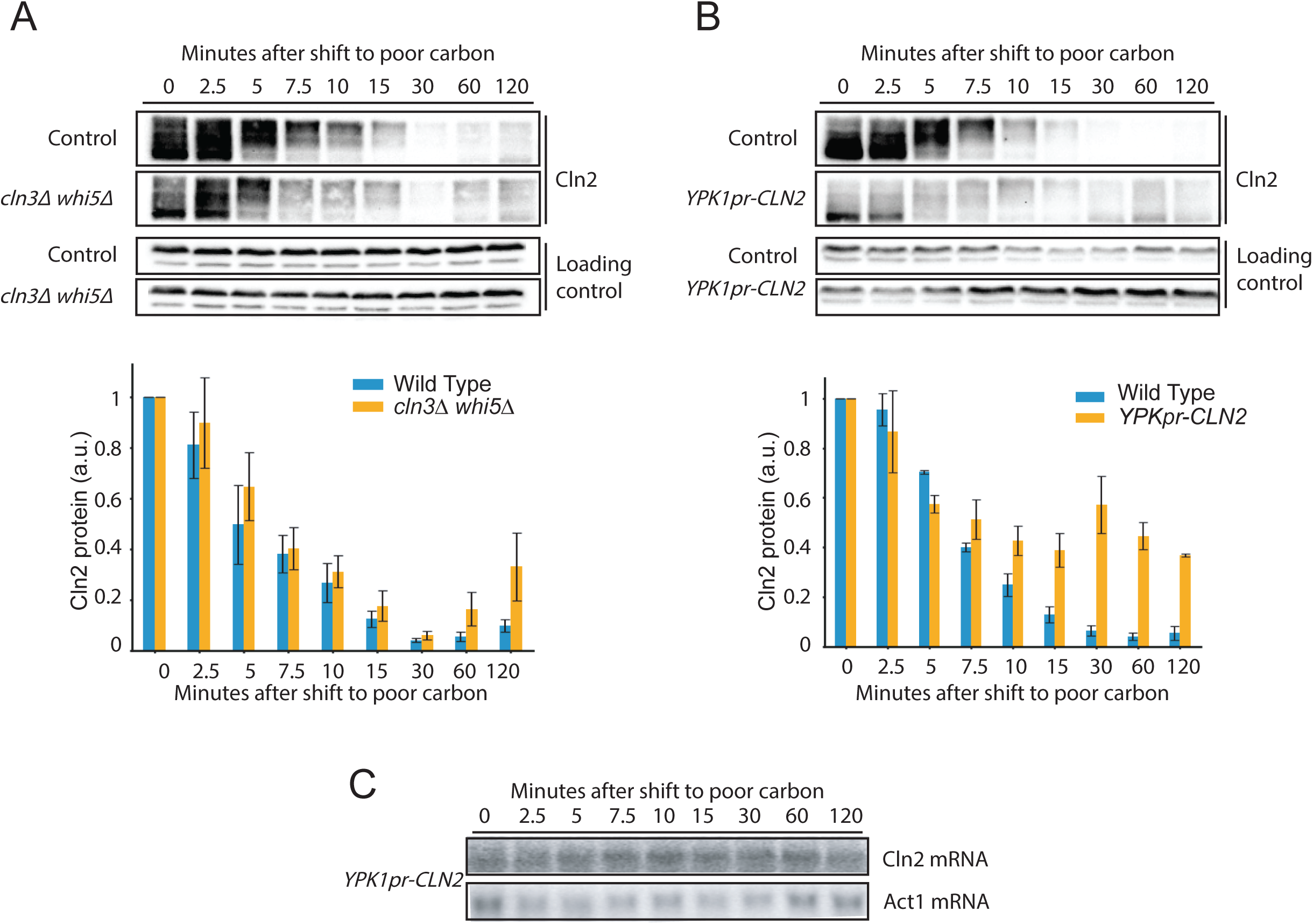
Nutrients modulate expression of Cln2 via post-transcriptional mechanisms. **(A)** Rapidly growing wild type or *cln3Δ whi5Δ* cells growing in rich carbon (YPD) were shifted to poor carbon (YPG/E) at 30°C,and the behavior of Cln2-9xMyc was analyzed by western blot. An anti-Nap1 antibody was used as a loading control. **(B)** Rapidly growing wild type or *YPK1pr-CLN2* cells growing in rich carbon (YPD) were shifted to poor carbon (YPG/E) at 30°C and the behavior of Cln2-9xMyc was analyzed by western blot. An anti-Nap1 antibody was used as a loading control.

To further test whether transcriptional mechanisms contribute to regulation of Cln2 levels, we replaced the *CLN2* promoter with the *YPK1* promoter, which is a constitutive promoter that does not show cell cycle dependent expression (Brambila et al., 2024). Cln2 protein expressed from the *YPK1* promoter (*YPK1pr-CLN2*) underwent hyperphosphorylation and destruction in response to a shift from rich to poor carbon, whereas Cln2 mRNA persisted (**Figures 3B and 3C**). Thus, nutrient modulation of Cln2 protein expression in response to a shift from rich to poor carbon occurs primarily at the post-transcriptional level. The kinetics of Cln2 protein destruction in *YPK1pr-CLN2* cells were slightly delayed relative to wild type. Previous studies found that Cln2 promotes its own transcription in a positive feedback loop, so the delay could be due to a loss of positive feedback. For example, in wild type cells loss of Cln2 protein would lead to loss of positive feedback, which would contribute to a rapid decline in Cln2 mRNA and protein. In the *YPK1pr-CLN2* cells, loss of Cln2 is likely driven entirely via destruction of the protein, which may take slightly longer.

We also tested the contribution of transcriptional mechanisms to control of Cln2 protein expression in cells synchronized in G1 phase. *YPK1pr-CLN2* cells were synchronized in early G1 phase by centrifugal elutriation and released into media containing rich or poor carbon, as in Figure 1. In both rich and poor carbon, Cln2 protein expressed from the *YPK1* promoter was repressed in early G1 phase and began to accumulate at the time of bud emergence after an interval of growth had occurred. Furthermore, the timing of Cln2 protein expression from the *YPK1* promoter was advanced and peak levels of Cln2 protein were reduced in poor carbon relative to rich carbon (**Figure 4**). Together, these results provide further evidence that post-transcriptional mechanisms play a major role in nutrient-dependent and cell cycle-dependent expression of Cln2 protein. A previous study found that cell cycle-dependent expression of Cln2 protein is also largely retained in *YPK1pr-CLN2* cells that were synchronized in G1 phase with mating pheromone (Brambila et al., 2024). There was some loss of repression of Cln2 in early G1 phase in the *YPK1pr-CLN2* cells, which indicates that transcriptional mechanisms help keep Cln2 protein levels low in early G1 phase. The primary function of transcriptional mechanisms may be to sharpen control of Cln2 protein expression. There is evidence that post-transcriptional mechanisms also play a major role in expression of mammalian G1 phase cyclins (Narasimha et al., 2014).

**Figure 4.**
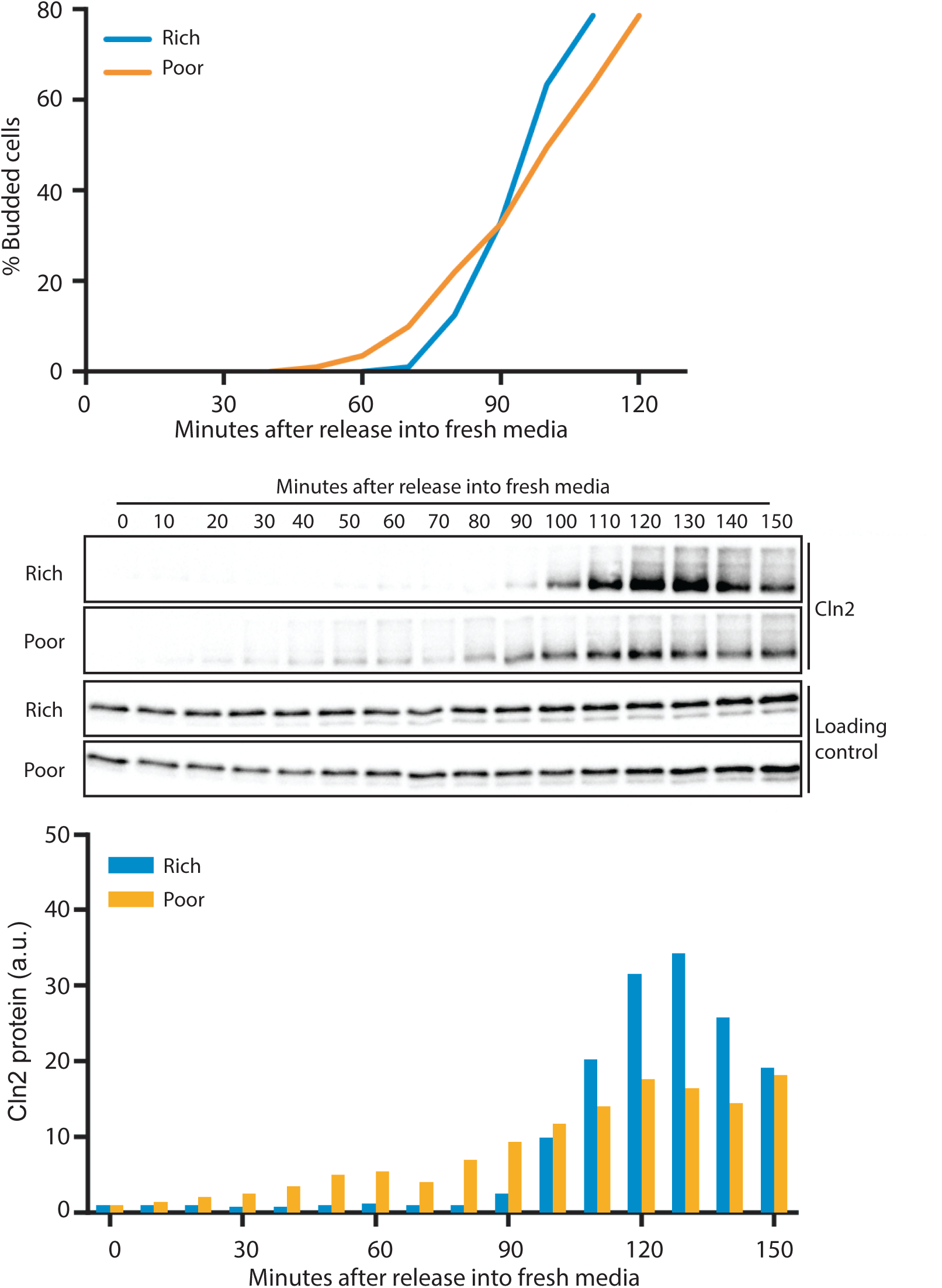
Expression of Cln2 is controlled by post-transcriptional mechanisms. *YPK1pr-CLN2* cells were grown to midlog phase in poor carbon (YPG/E) and small unbudded cells were isolated by centrifugal elutriation. Cells were released into either rich carbon (YPD) or poor carbon (YPG/E) medium at 25°C and samples were taken at 10 min intervals to assay bud emergence and Cln2-9xMyc protein expression. All data in the figure are from the same biological replicate. An anti-Nap1 antibody was used as a loading control.

### Phosphorylation of Cln2 is required for nutrient modulation of Cln2 protein levels

The decrease in Cln2 levels that occurs after a shift from rich to poor carbon is preceded by hyperphosphorylation of Cln2. We therefore hypothesized that phosphorylation of Cln2 targets it for increased turnover in poor carbon. Previous work found that Cln2 protein turnover is controlled by a phosphodegron (Lanker et al., 1996; Willems et al., 1996). Mutation of the 7 phosphorylation sites in the degron to alanines causes a 4-fold increase in Cln2 protein levels and a loss of cell cycle-dependent regulation of Cln2 protein expression. It has generally been assumed that the phospho-degron serves to target Cln2 for destruction at the end of G1 phase when it is no longer needed, but no previous experiments have ruled out the possibility that the phospho-degron has other functions. To test whether the degron is required for nutrient modulation of Cln2 protein levels, we used an allele of *CLN2* in which the phospho-degron sites have been converted to alanines (*cln2-4T3S*). Wild type and *cln2-4T3S* cells were shifted from rich to poor carbon and Cln2 protein levels and phosphorylation were assayed by western blot (**Figure 5**). The cln2-4T3S protein failed to undergo phosphorylation and to show a decrease in protein levels after a shift to poor nutrients. These results show that nutrients control Cln2 protein levels via a phospho-degron.

**Figure 5.**
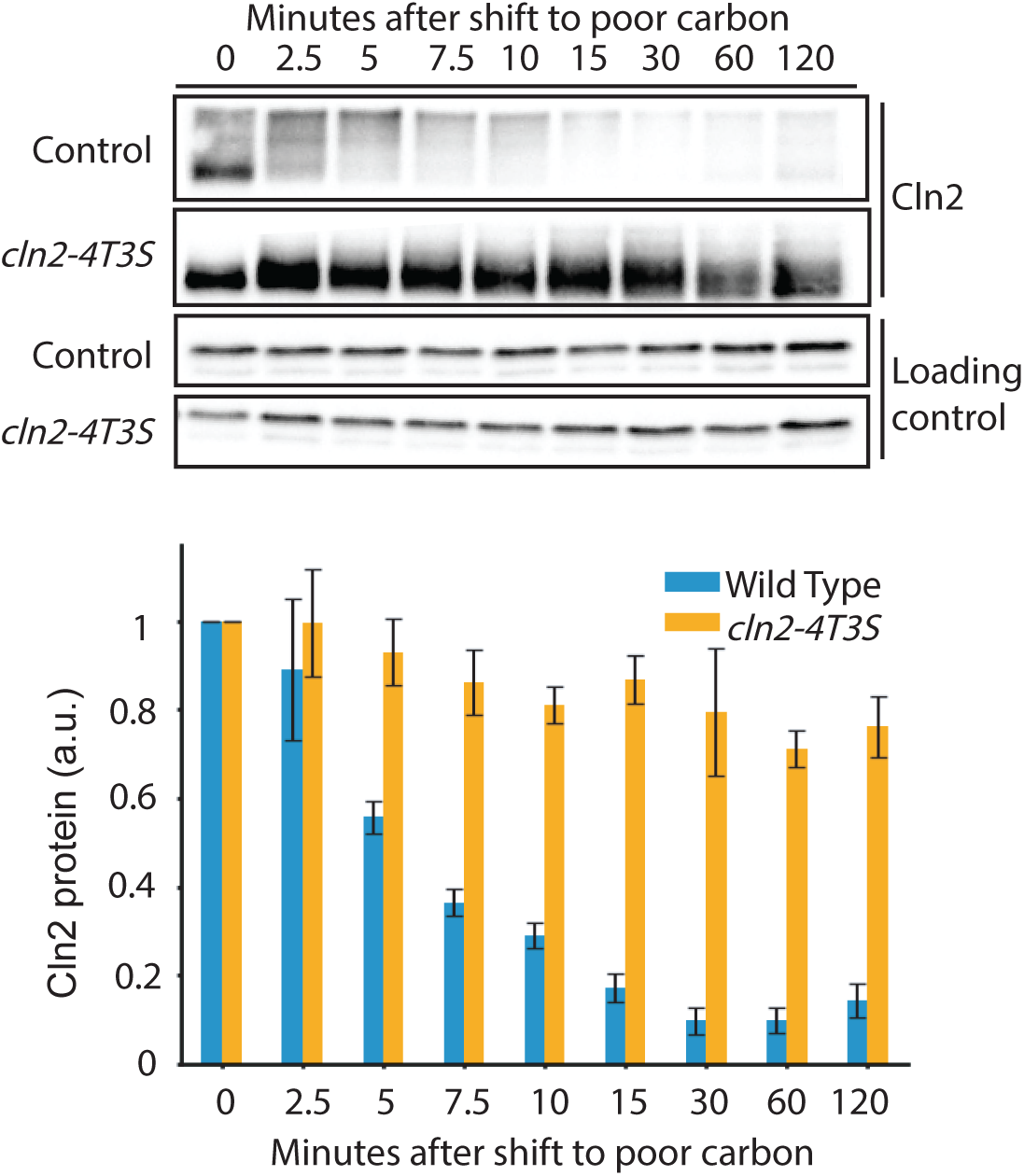
Phosphorylation of Cln2 is required for nutrient modulation of Cln2 protein levels. Rapidly growing *cln2-4T3S-9XMyc* cells in rich carbon (YPD) were shifted to poor carbon (YPG/E) at 25°C and the behavior of *cln2-4T3S-9XMyc* was analyzed by western blot. An anti-Nap1 antibody was used as a loading control.

### PP2A^Cdc55^ is required for normal control of Cln2 protein levels and phosphorylation, but is not required for nutrient modulation of Cln2 protein levels

Phosphorylation and destruction of Cln2 in response to a shift to poor carbon could be initiated by activation of a kinase, inhibition of a phosphatase, or both. We first searched for a phosphatase that could play a role in controlling phosphorylation of Cln2 in response to a shift from rich to poor carbon. Previous work suggested that PP2A^Cdc55^ controls Cln2 phosphorylation (McCourt et al., 2013). PP2A^Cdc55^ is a highly conserved trimeric phosphatase complex that includes a catalytic subunit, a scaffolding subunit, and Cdc55, which is thought to be a targeting subunit that confers specificity. Cells that lack Cdc55 are viable but show slow proliferation as well as defects in cell size and morphology that indicate aberrant control of cell growth (Healy et al., 1991; Yang et al., 2000; Pal et al., 2008). Previous work found that Cln2 levels are substantially lower in *cdc55Δ* cells but did not achieve clear resolution of phosphorylated forms of Cln2 and did not test whether Cdc55 plays a role in nutrient-dependent control of Cln2 phosphorylation (McCourt et al., 2013). We found that Cln2 protein was hyperphosphorylated and protein levels were reduced in *cdc55Δ* cells (**Figure 6A**). The fact that Cln2 is not quantitatively shifted to the fully phosphorylated form in *cdc55Δ* cells could indicate that there is another phosphatase that opposes phosphorylation of Cln2. Alternatively, the unphosphorylated Cln2 that is detected in *cdc55Δ* cells could be a fraction of Cln2 that has not yet experienced phosphorylation by a protein kinase. When *cdc55Δ* cells were shifted from rich to poor carbon, Cln2 underwent phosphorylation and destruction, which indicates that hyperphosphorylation of Cln2 in this context is not due simply to inhibition of PP2A^Cdc55^ (**Figure 6B)**. Overall, the results are most consistent with a model in which a shift from rich to poor carbon activates a kinase that phosphorylates Cln2, but do not rule out the possibility that a shift to poor carbon inhibits a phosphatase that is not PP2A^Cdc55^.

**Figure 6.**
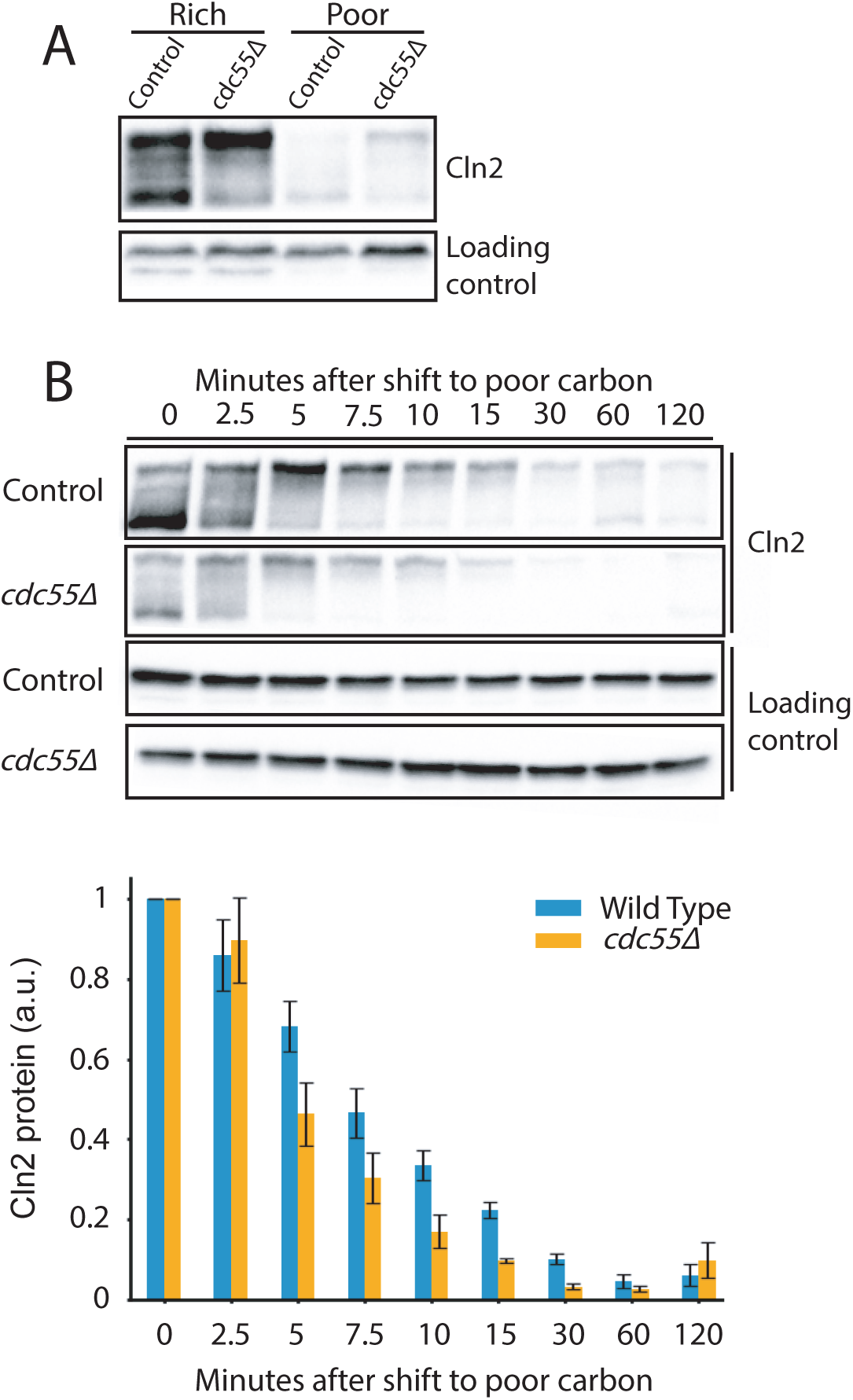
PP2A^Cdc55^ is required for normal control of Cln2 protein phosphorylation and expression but is not required for nutrient modulation of Cln2 protein levels. **(A)** Wild type and *cdc55Δ* cells were grown to log phase in rich carbon (YPD) or poor carbon (YPG/E) medium and levels of Cln2-9XMyc were analyzed by western blot. An anti-Nap1 antibody was used as a loading control. **(B)** Rapidly growing wild type or *cdc55Δ* cells in rich carbon (YPD) were shifted to poor carbon (YPG/E) at 30°C and the behavior of Cln2-9xMyc was analyzed by western blot. An anti-Nap1 antibody was used as a loading control.

### Cdk1 activity is not required for phosphorylation and destruction of Cln2 in response to poor carbon

We next search for kinases that promote phosphorylation of Cln2. Previous work reached the conclusion that Cdk1 is responsible for phosphorylation of the Cln2 phospho-degron, likely via autophosphorylation events (Lanker et al., 1996). However, these studies relied on a temperature sensitive allele of Cdk1 and prolonged incubation of cells at the restrictive temperature (3 hours), which could cause indirect effects on Cln2 phosphorylation. We used an analog-sensitive allele of Cdk1 (*cdk1-as*) to test whether Cdk1 activity is required for phosphorylation and destruction of Cln2 in response to a shift to poor nutrients. Wild type and *cdk1-as* cells were shifted from rich to poor nutrients in the presence of 1NM-PP1. Inhibition of Cdk1 had only a modest effect on phosphorylation and destruction of Cln2 (**Figure 7**). There was an overall reduction of Cln2 phosphorylation in *cdk1-as* cells, which suggests that Cdk1 activity contributes to Cln2 phosphorylation, but the data suggest that Cdk1 activity does not make a substantial contribution to nutrient modulation of Cln2 protein.

**Figure 7.**
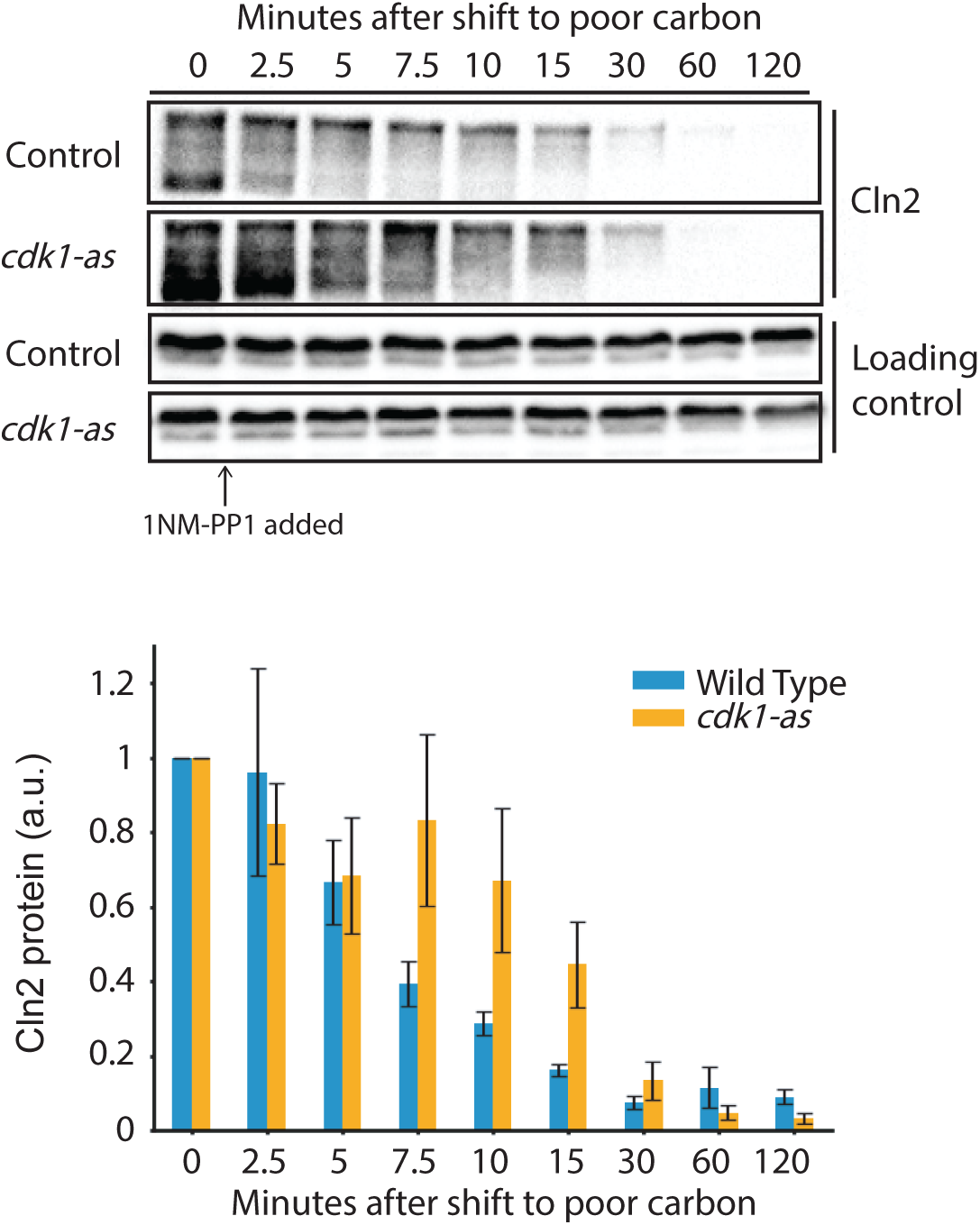
Cdk1 activity is not required for phosphorylation and destruction of Cln2 in response to poor carbon. Rapidly growing wild type or *cdk1-as* cells in rich carbon (YPD) were shifted to poor carbon (YPG/E) medium containing 5µM 1NM-PP1 at 30°C and the behavior of Cln2-9XMyc was analyzed by western blot. An anti-Nap1 antibody was used as a loading control.

### A MAP kinase cascade is required for phosphorylation and destruction of Cln2 in response to poor carbon

The phosphorylation sites within the degron that targets Cln2 for destruction are all serines or threonines followed by a proline, which indicates that the relevant kinase is a proline-directed kinase. Beyond Cdks, another important family of proline-directed kinases is MAP kinases. Previous work suggested that transcription of the *CLN2* gene is controlled by a MAP kinase cascade (Madden et al., 1997; Baetz et al., 2001; Gray et al., 1997). In this cascade, the MAPKKK is Bck1 and the MAPKK is encoded by a pair of redundant paralogs referred to as Mkk1 and Mkk2. The MAPK is encoded by a pair of paralogs referred to as Slt2 and Kdx1, with Slt2 playing a dominant role in the pathway. Complete loss of function of components of the cascade causes slow proliferation and a range of defects, including defects in control of cell growth, as cells become abnormally large in rich nutrient conditions or abnormally small in poor nutrient conditions (Schneider et al., 2004; Mazzoni et al., 1993). Cells that lack the pathway also show a tendency to undergo lysis. No previous studies have tested whether the MAP kinase cascade controls Cln2 protein expression via post-transcriptional mechanisms. Since Cln2/Cdk1 influences *CLN2* transcription via a positive feedback loop, the previously observed effects of the MAP kinase cascade on CLN2 transcription could work through the feedback loop.

We used *bck1Δ* cells to test whether the MAP kinase cascade is required for nutrient modulation of Cln2 protein levels. Wild type and *bck1Δ* cells growing asynchronously in rich carbon were rapidly shifted to poor carbon and levels of Cln2 protein and Cln2 phosphorylation were assayed by western blot (**Figure 8A**). Loss of Bck1 caused a failure in nutrient-dependent phosphorylation and destruction of Cln2. Before the shift to poor carbon, phosphorylated forms of Cln2 can be detected in the *bck1Δ* cells, which indicates that the MAP kinase cascade is not solely responsible for phosphorylation of Cln2. We also assayed activity of the MAP kinase cascade using a phospho-specific antibody that detects the active form of Slt2 (**Figure 8B**). MAP kinase activity showed a transient decrease after the shift to poor carbon and then underwent an increase that was correlated with hyperphosphorylation of Cln2. The active form of Slt2 is undetectable in *bck1Δ* cells, which confirmed that the MAP kinase pathway is fully inactivated.

**Figure 8.**
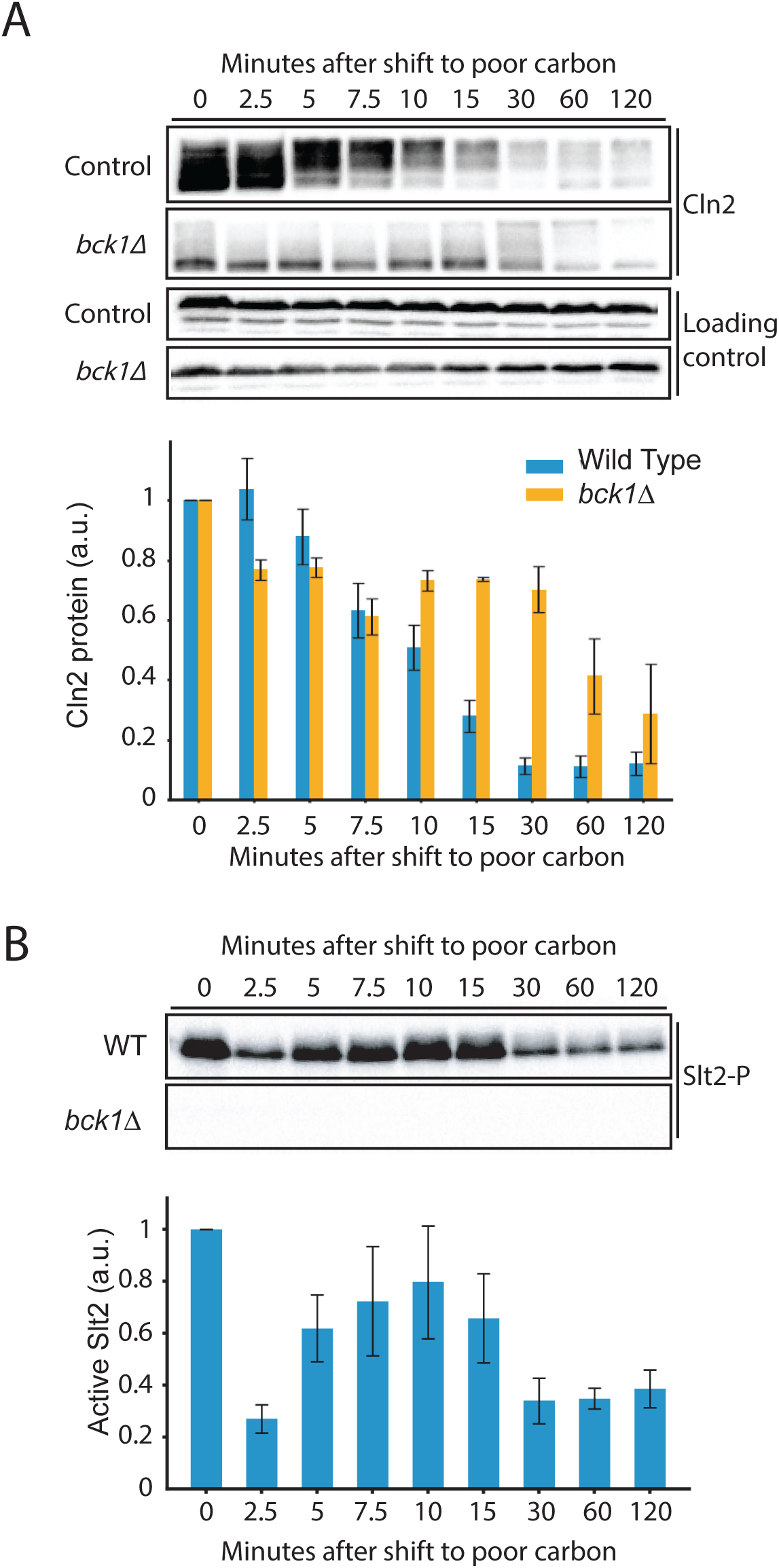
A MAP kinase cascade is required for phosphorylation and destruction of Cln2 in response to poor carbon. **(A)** Rapidly growing wild type or *bck1Δ* cells in rich carbon (YPD) were shifted to poor carbon (YPG/E) medium at 30°C and the behavior of Cln2-9xMyc was analyzed by western blot. An anti-Nap1 antibody was used as a loading control. **(B)** Levels of the active form of Slt1 MAP kinase were assayed by western blot using a phospho-specific antibody that recognizes the active form of Slt2.

There are several models that could explain these results. In one model, a shift to poor carbon activates Slt2 to drive phosphorylation of Cln2, either directly or indirectly via activation of another kinase. Since a substantial amount of the active form of Slt2 is already present in rich carbon before the shift, this model would suggest that a shift to poor carbon redirects Slt2 activity to more efficiently promote phosphorylation of Cln2, potentially via changes in localization. An alternative model is that Slt2 inhibits a phosphatase that opposes phosphorylation of Cln2. The models are not mutually exclusive, as the MAP kinase cascade could drive both phosphorylation of Cln2 and inhibition of an opposing phosphatase.

### TORC2 signaling is required for phosphorylation and destruction of Cln2 in response to poor carbon

Genetic data suggest that the Slt2 MAP kinase cascade is controlled by TOR complex 2 (TORC2) and the protein kinase Pkc1 (Helliwell et al., 1998a; b; Schmidt et al., 1997; Nonaka et al., 1995; Irie et al., 1993; Lee and Levin, 1992; Lee et al., 1993). For example, *pkc1* mutants are suppressed by gain-of-function alleles of *BCK1* and *MKK1*, and gain-of-function alleles of *PKC1* suppress TORC2 mutants. Pkc1 is most closely related to mammalian PKN kinases and is required for viability. Previous work suggested that Pkc1 plays roles in control of bud growth and Cln2 transcription (Gray et al., 1997; Anastasia et al., 2012; Madden et al., 1997).

To test whether TORC2 plays a role in activation of the MAP kinase cascade, we added AID tags to two essential subunits of TORC2 (Avo1 and Avo3). Use of an AID tag to inactivate Avo1 or Avo3 alone caused slow proliferation but did not cause lethality, likely because AID tags do not fully deplete proteins. Inactivation of Avo1 and Avo3 together caused a stronger effect on proliferation (**Figure S1**). Avo1/3-AID cells were grown to log phase in rich YPD medium. Auxin was then added for 30 minutes to degrade Avo1 and Avo3 and the cells were rapidly washed into poor carbon YPG/E medium containing auxin and Cln2 protein was monitored by western blot. We also monitored levels of the active form of Slt2. Hyperphosphorylation of Cln2 failed to occur in the Avo1/3-AID cells and the active form of Slt2 disappeared (**Figure 9A**). These results are consistent with a model in which TORC2-dependent signals control Cln2 protein levels via the MAP kinase cascade.

**Figure 9.**
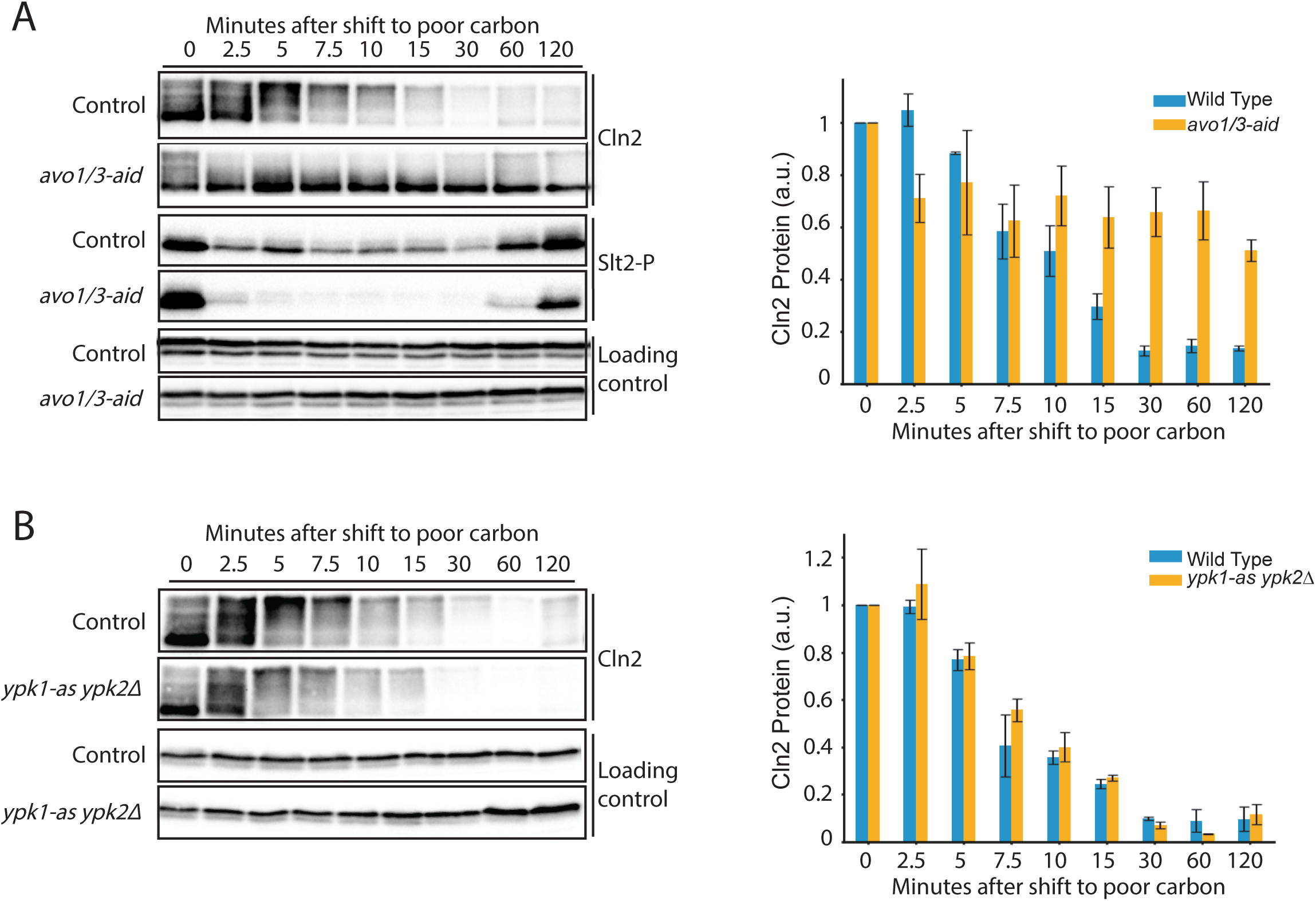
TORC2 signaling is required for phosphorylation and destruction of Cln2 in response to poor carbon. **(A)** Rapidly growing wild type or *avo1/3-AID* cells in rich carbon (YPD) were shifted to poor carbon (YPG/E) medium containing 1 mM auxin at 30°C and the behavior of Cln2-9xMyc was analyzed by western blot. An anti-Nap1 antibody was used as a loading control. **(B)** Rapidly growing wild type or *ypk1-as ypk2Δ* cells in rich carbon (YPD) were shifted to poor carbon (YPG/E) medium containing 10 uM 3-MOB-PP1 at 30°C and the behavior of Cln2-9xMyc was analyzed by western blot. An anti-Nap1 antibody was used as a loading control.

TORC2 also signals via the kinases Ypk1 and Ypk2, which are homologs of mammalian SGK kinases (Kamada et al., 2005; Niles et al., 2012). Ypk1 and Ypk2 are redundant paralogs. Cells that lack either paralog are viable, while loss of both is lethal. We found that inhibition of an analog-sensitive allele of Ypk1 in a *ypk2Δ* background (*ypk1-as ypk2Δ*) had no effect on hyperphosphorylation and destruction of Cln2 in response to a shift from rich to poor carbon (**Figure 9B**). Thus, TORC2 appears to influence Cln2 phosphorylation primarily via a Pkc1-MAP kinase axis.

Previous work has shown that the TORC2 network is required for modulation of growth rate and cell size in response to changes in carbon source (Lucena et al., 2018). The finding that TORC2 signaling is required for modulation of Cln2 protein levels is therefore consistent with a model in which nutrients modulate Cln2 protein levels to match growth rate to nutrient availability in G1 phase.

## Discussion

Expression of late G1 phase cyclins is the critical molecular event that marks commitment to enter the cell cycle. In budding yeast, the late G1 phase cyclins drive growth of a new daughter bud (Cross, 1990; McCusker et al., 2007). Nutrients strongly influence cell cycle entry, expression of late G1 phase cyclins, and the duration and extent of growth in G1 phase. Here, we investigated signals that control expression of late G1 phase cyclins, with the goal of better understanding their roles and regulation, and how nutrient availability influences their expression and function.

### Modulation of Cln2 protein expression in response to nutrient availability occurs via post-transcriptional mechanisms

In previous work we found evidence that transcriptional mechanisms play a relatively minor role in control of Cln2 protein expression (Brambila et al., 2024). Thus, cells that lack both Cln3 and Whi5, which are thought to play a major role in control of *CLN2* transcription, show normal cell cycle-dependent expression of Cln2 protein. Cells in which the *CLN2* promoter has been replaced with a heterologous constitutive promoter also retain substantial cell cycle-dependent expression of Cln2, and neither Whi5 nor the *CLN2* promoter are required for the ability of Cln3 to influence Cln2 protein expression. Here, we extended these studies by analyzing Cln2 protein expression in cells synchronized via centrifugal elutriation, which is a minimally disruptive procedure that avoids any potential artifacts that may be due to synchronization with mating pheromone. We again found that Cln2 expressed from a heterologous constitutive promoter shows nearly normal expression – Cln2 protein expression was suppressed in early G1 phase and appeared only after an interval of growth in early G1 phase. Thus, post-transcriptional mechanisms play a major role growth-dependent and cell cycle-dependent expression of Cln2 protein. We also investigated how nutrient availability influences expression of Cln2. Previous studies using asynchronous cells found that Cln2 protein expression is reduced in poor nutrients (Schneider et al., 2004). This has always been a puzzling observation, as it has been unclear why the expression and requirement for G1 cyclins should be reduced in poor nutrients. By identifying the signals that influence late G1 phase cyclin expression in response to nutrient availability we aimed to learn more about how and why cyclin protein levels are reduced in poor nutrients. We found that shifting asynchronously growing cells from rich to poor carbon causes rapid hyperphosphorylation and destruction of Cln2. This discovery shows that expression of Cln2 is tightly linked to signals that monitor the availability of nutrients that are required for cell growth. In synchronized cells, we found that growth in poor carbon causes entry into the cell cycle at a smaller cell size and with lower levels of Cln2. These data confirm a previous study in asynchronous cells that suggested that cells enter the cell cycle with reduced levels of Cln2 in poor nutrients (Schneider et al., 2004). Together, the data indicate that Cln2 protein expression is correlated with the rate of growth set by nutrient availability. The data are also consistent with the possibility that Cln2 protein levels are correlated with the extent of growth. For example, Cln2 could first be detected when cells were at approximately 5% bud emergence and protein levels then increased steadily. One interpretation of this observation is that Cln2 first appears when bud growth is initiated and then accumulates to higher levels as the bud grows, which would suggest that expression of Cln2 is correlated with the extent of bud growth. However, the limited synchrony of the cell population makes it difficult to rule out alternative explanations.

Nutrient-dependent modulation of Cln2 protein expression occurred normally in cells that lack Cln3 and Whi5 and in cells in which the normal Cln2 promoter was replaced with a heterologous constitutive promoter. Thus, nutrient-dependent control of Cln2 protein expression occurs primarily via post-transcriptional mechanisms.

### A TORC2-Pkc1-MAP kinase signaling axis is required for nutrient modulation of Cln2 expression

We found that hyperphosphorylation of Cln2 in response to poor carbon requires a previously identified phospho-degron that was thought to work primarily at the end of G1 phase to destroy Cln2 when it is no longer needed. Our results indicate that the phospho-degron plays more diverse roles in control of Cln2 protein expression.

A candidate approach led to the discovery that a MAP kinase cascade is required for phosphorylation and destruction of Cln2 in response to poor carbon. We also found that TORC2 signaling is required both for normal nutrient-dependent regulation of the MAP kinase cascade and for nutrient modulation of Cln2 protein expression. Previous studies suggest that TORC2 directly activates Pkc1, and that Pkc1 controls the MAP kinase cascade (Helliwell et al., 1998a; b; Schmidt et al., 1997; Nonaka et al., 1995; Irie et al., 1993; Lee and Levin, 1992; Lee et al., 1993). The dynamics of MAP kinase signaling in this context appear to be complex. For example, substantial amounts of the active form of Slt2 kinase can be detected in cells growing in rich carbon with high levels of Cln2, and a fraction of Cln2 is hyperphosphorylated. A shift to poor carbon causes a transient decrease in the amount of active Slt2, followed by an increase in active Slt2 that is correlated with hyperphosphorylation and destruction of Cln2. The amount of active Slt2 detected during hyperphosphorylation of Cln2 in poor carbon is lower than the amount detected in rich carbon, so why is Cln2 hyperphosphorylated and destroyed in poor carbon but not in rich carbon? A model that could explain these data is that a shift to poor carbon redirects the activity of Slt2 to Cln2, potentially via a change in localization. The shift to poor carbon could also inhibit a phosphatase that opposes phosphorylation of Cln2. The data do not yet distinguish whether Cln2 is directly phosphorylated by MAP kinase, or whether MAP kinase activates another kinase to phosphorylate Cln2. Inhibition of Cdk1 had little effect on hyperphosphorylation of Cln2 in response to poor carbon, so Cdk1 is unlikely to play an important role.

There is evidence that Pkc1 functions in a mechanism that measures bud growth during the initial polar phase of growth that is driven by Cln2/Cdk1 activity (Anastasia et al., 2012). Pkc1 is localized to the tip of the growing bud and undergoes gradual multisite phosphorylation during bud growth. Phosphorylation of Pkc1 is dependent upon membrane trafficking events that drive plasma membrane growth in the growing bud, and the extent of phosphorylation is correlated with the extent of growth. Once polar bud growth is complete, Pkc1 drives removal of Cdk1 inhibitory phosphorylation to initiate the transition into mitosis and into the next growth phase that occurs throughout mitosis. It is unknown whether the fraction of Pkc1 that undergoes growth-dependent phosphorylation also regulates the MAP kinase cascade.

Together, the data suggest a model in which nutrients modulate Cln2 protein expression via a TORC2-Pkc1-MAP kinase signaling axis. Since TORC2 is closely associated with regulation of cell growth and size (Szwed et al., 2021; Emmerstorfer-Augustin and Thorner, 2023; Lucena et al., 2018), the data further suggest that regulation of Cln2 is closely associated with control of cell growth.

### G1 phase cyclins are closely associated with cell growth

Our analysis of Cln2 expression, combined with previous analysis of Cln3 expression, indicate that both early and late G1 phase cyclins have a unique association with cell growth. Cln3 protein levels increase gradually during growth in G1 phase and are correlated with the rate and extent of growth (Sommer et al., 2021). Our analysis of Cln2 protein expression is consistent with the possibility that Cln2 protein levels are similarly correlated with the rate and extent bud growth. A previous study also found evidence that Cln2 protein levels are set to match the growth rate set by nutrient availability (Schneider et al., 2004). Levels of both Cln3 and Cln2 are rapidly down-regulated in response to a shift from rich to poor nutrients that leads to a rapid decrease in growth rate, and levels of both proteins are controlled by TORC2-dependent signals (Sommer et al., 2021). Cln2 shows an especially close association with cell growth because it is required for initiation of bud growth and Cln2/Cdk1 activity is continuously required for the initial polar growth phase of the bud (McCusker et al., 2007). No previous analysis has tested whether Cln3 is required for normal cell growth. *cln3Δ* cells are abnormally large, but this doesn’t rule out the possibility that loss of Cln3 causes slow growth. Testing whether Cln3 is required for cell growth is difficult because *cln3Δ* cells have severe size defects that make it difficult to determine the immediate consequences of loss of Cln3 function, and there are no good tools available that would allow rapid conditional inactivation of Cln3 in early G1 phase. Expression of the mitotic cyclin Clb2 is not modulated by nutrient availability (Figure 2C) and mitotic cyclins are not required for bud growth (Fitch et al., 1992), which provides further evidence that G1 phase cyclins have a unique association with cell growth.

The close association between Cln2 and cell growth suggests a potential explanation for the mysterious down-regulation of G1 phase cyclins in poor nutrients. Since Cln2/Cdk1 activity is continuously required for bud growth (McCusker et al., 2012, 2007), the decreased expression of Cln2 protein in poor nutrients could lead to a slower rate of bud growth that matches growth rate to nutrient availability. Future experiments will test this model.

## Materials and Methods

### Yeast strains, media, plasmids, and inhibitors

All strains are in the W303 background (leu2-3, 112 ura3-1 can1-100 ade2-1 his3-11,15 trp1-1 GAL+ ssd1-d2 bar1) and derived from the parent strain DK186. Table 1 shows additional genetic features. One-step, PCR-based gene replacement was used for making deletions and adding epitope tags at the endogenous locus (Longtine et al., 2000; Janke et al., 2004).

**Table 1:** Strains with multiple strain numbers indicate multiple independent isolates of the same strain.

| Strain | Genotype | Reference or Source | Figures |
| --- | --- | --- | --- |
| DK186 | *MATa* | Altman and Kellogg, 1997 | N/A (background strain) |
| DK4582 | *CLN2-9MYC::hphNT1* | This study | Figures 1 and 2 |
| DK4714 | *His3-YPK1-CLN2-9MYC::hphNT1* | Brambila et al., 2024 | Figure 3B and C<br>Figure 4 |
| DK4815 | *CLN2-9myc::clonat*<br>*whi5Δ::KANMX4*<br>*cln3Δ::His3* | This study | Figure 3A |
| DK4780 | *CLN2(4T3S)-9MYC::hphNT1* | This study | Figure 5 |
| DK4733 | *CLN2-9MYC::hphNT1*<br>*cdc55Δ::KanMx4* |  | Figure 6 |
| DK4722 | *cdc28::cdc28-as1*<br>*CLN2-9Myc::hphNT1* | This study | Figure 7 |
| DK4935 | *CLN2-9MYC::hphNT1*<br>*bck1Δ::KanMX4* | This study | Figure 8 |
| DK4855 | *his3::HIS3+TIR1*<br>*leu2::LEU2+TIR1* | This study | Figure 9A |
| DK5054 | *his3::HIS3+TIR1*<br>*leu2::LEU2+TIR1*<br>*avo1-miniAID::clonat* | This study | N/A |
| DK4856 | *his3::HIS3+TIR1*<br>*leu2::LEU2+TIR1*<br>*avo3-AID::KanMX* | This study | N/A |
| DK5025 | *his3::HIS3+TIR1*<br>*leu2::LEU2+TIR1*<br>*CLN2-9MYC::hphNT1*<br>*avo3-AID::KanMX*<br>*avo1-miniAID::clonat* | This study | Figure 9A |
| DK5067 | *Ypk2Δ::His3*<br>*Ypk1-as (L424A)*<br>*CLN2-9MYC::hphNT1* | This study | Figure 9B |

Cells were grown in YP medium (1% yeast extract, 2% peptone) that contained 40 mg/L supplemental adenine and a carbon source, except where noted in the figure legends. Rich carbon medium (YPD) contained 2% dextrose, while poor carbon medium (YPG/E) contained 2% glycerol and 2% ethanol. In experiments using the ATP analog inhibitor 3-MOB-PP1, no additional adenine was added to the media.

Stock solutions of ATP analog inhibitors were solubilized in 100% DMSO at 10 mM. 3-MOB-PP1 was a gift from Joshua Combs, Jack Stevenson and Kevan Shokat (UCSF). Auxin (in-dole-3-acetic acid) (Aldrich) stock was prepared at 50 mM in 100% EtOH and used at 1 mM.

### Centrifugal elutriation

Strains were grown in YPG/E medium overnight to an OD600 of 0.4–0.8 at 30°C. Cells were harvested by centrifugation at 4000 rpm in a JLA 8.1 rotor at 4°C for 6 min. Cell pellets were resuspended in ∼100 mL cold YPG/E and then sonicated for 1 min at duty cycle 0.5 using a Braun-Sonic U sonicator with a Braun 2000U probe at 4°C. Cells were loaded onto a Beckman JE-5.0 elutriator in a Beckman Coulter J6-MI centrifuge at 2900 rpm at 4°C. After all cells were loaded, fluid flow was continued for 10 min to allow equilibration. Pump speed was then increased gradually to collect small unbudded cells, which were collected and pelleted by spinning in a JA-14 rotor at 5000 rpm for 5 min. Cells were resuspended into fresh medium at OD600 0.4–0.6 and grown at 25°C in a shaking water bath. For western blotting, 1.6 mL samples were collected at each time point as cells progressed through G1 phase. Since the cells do not divide during the time course, a constant number of cells is collected at each time point so that signals are normalized to cell number. Thus, western blot signals represent protein copy number per cell, rather than protein concentration. For Coulter counter analysis, samples were collected and fixed with 3.7% formaldehyde for 30 min for cell size measurements and to calculate the percentage of budded cells. Median cell size was calculated by the Coulter AccuComp software for each time point to plot cell size during growth in G1.

### Western blotting

1.6 mL samples of cells taken from cultures were pelleted in a microfuge at 13,200 rpm for 15 s before aspirating the supernatant and adding 250 μL of glass beads and freezing on liquid nitrogen. Cells were lysed in 140 μL of 1× SDS-PAGE sample buffer (65 mM Tris-HCl, pH 6.8, 3% SDS, 10% glycerol, 100 mM **β**-glycerophosphate, 50 mM NaF, 5% **β**-mercaptoethanol, 2 mM PMSF, and bromophenol blue) by bead beating in a Biospec Mini-Beadbeater-16 at 4°C for 2 min. The lysate was centrifuged for 15 s to bring the sample to the bottom of the tube and was then incubated in a 100°C water bath for 5 min followed by centrifugation for 5 min at 13,200 rpm. Depending on the protein blot, 10–15 μL of the lysate were loaded into 10% acrylamide SDS-PAGE gels, then run at a constant current setting of 20 mA per gel at a maximum of 165 V. Gels were transferred to nitrocellulose membrane in a Bio-Rad Trans-Blot Turbo Transfer system. Blots were probed overnight at 4°C in 4% milk in western wash buffer (1× PBS + 250 mM NaCl +0.1% Tween-20) with mouse monoclonal Myc Tag (9B11) mouse antibody (Cell Signaling Technology, #2276S), polyclonal anti-Nap1 antibody, polyclonal Phospho-p44/42 MAPK antibody (Cell Signaling Technology #4370), or polyclonal rabbit anti-T662P antibody (gift from Ted Powers, University of California, Davis). Western blots using anti-T662P antibody were first blocked using TBST (10 mM Tris-Cl, pH 7.5, 100 mM NaCl, and 0.1% Tween-20) + 4% milk, followed by one wash with TBST, then incubated overnight with anti-T662P antibody in TBST +4% BSA. Western blots were incubated in secondary donkey antimouse (GE Healthcare NA934V) or donkey antirabbit (GE Healthcare NXA931 or Jackson Immunoresearch 711-035-152) antibody conjugated to HRP at room temperature for 60–90 min before imaging with Advansta ECL chemiluminescence reagents in a Bio-Rad ChemiDoc imaging system.

### Log phase nutrient shift

Cells were grown to midlog phase (OD600 0.4–0.7) in rich carbon (YPD) at 25°C. Cells were shifted to poor carbon by pelleting the cells and resuspending in YPG/E medium two times at room temperature grown and then incubated with gentle shaking at 25°C or 30°C (see figure legends)

### Western blot quantification

Western blots were quantified using Bio-Rad Imagelab software v6.0.1. For the elutriation and time course experiments, relative signal was calculated as a ratio of the raw signal at each time point over the raw signal at the zero time point.

### Northern blotting

Gel-purified PCR products were used to generate 32P-labeled probes to detect CLN2 and ACT1 mRNAs by northern blotting (CLN2 oligonucleotides: TATTACTTGGG-TATTGCCCATACCAAAAAGA, TGAACCAATGATCAATGATTACGT; ACT1 oligonucleotides: TCATACCTTCTACAACGAATTGAGA and ACACTTCATGATGGAGTTGTAAGT). Northern blotting was carried out as previously described (Cross and Tinkelenberg, 1991; Kellogg and Murray, 1995).

### Experimental replicates

All experiments were repeated for a minimum of three biological replicates. Biological replicates are defined as experiments carried out on different days with different starting cultures.

## Acknowledgements

We thank members of the lab for helpful discussions, and Joshua Combs, Jack Stevenson and Kevan Shokat for providing 3-MOB-PP1. This work was funded by NIH grant R35 GM131826.

## Author contributions

Navid Zebarjadi, Formal analysis, Investigation, Methodology, Writing - original draft, Writing review and editing; Douglas R Kellogg, Conceptualization, Funding acquisition, Investigation, Methodology, Project administration, Writing - original draft, Writing - review and editing

---

**Figure S1.**
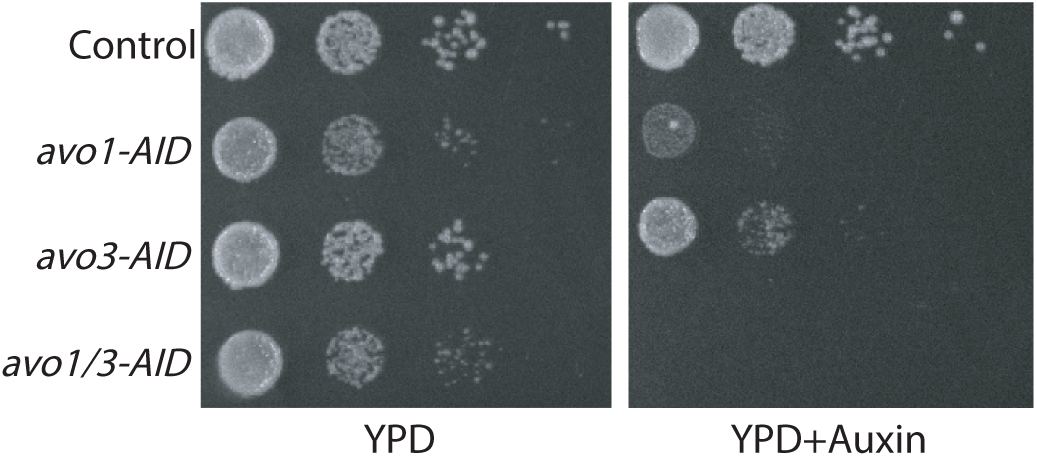
Growth of AID-tagged alleles of AVO1 and AVO3 in the presence and absence of 1 mM auxin. Serial 10-fold dilutions of cells of the indicated genotypes were spotted onto YPD or YPD + 1 mM auxin and grown at 30°C.

